# Role of Ethanolamine Utilization and Bacterial Microcompartment Formation in *Listeria monocytogenes* Intracellular Infection

**DOI:** 10.1101/2023.12.19.572424

**Authors:** Ayan Chatterjee, Karan Gautam Kaval, Danielle A. Garsin

## Abstract

Ethanolamine (EA) affects the colonization and pathogenicity of certain human bacterial pathogens in the gastrointestinal tract. However, EA can also affect the intracellular survival and replication of host-cell invasive bacteria such as *Listeria monocytogenes* (LMO) and *Salmonella enterica* serovar Typhimurium (*S.* Typhimurium). The EA utilization (*eut)* genes can be categorized as regulatory, enzymatic, or structural, and previous work in LMO showed that loss of genes encoding functions for the enzymatic breakdown of EA inhibited LMO intracellular replication. In this work, we sought to further characterize the role of EA utilization during LMO infection of host cells. Unlike what was previously observed for *S.* Typhimurium, in LMO, an EA regulator mutant (*ΔeutV)* was equally deficient in intracellular replication compared to an EA metabolism mutant (*ΔeutB*), and this was consistent across Caco-2, RAW 264.7 and THP-1 cell lines. The structural genes encode proteins that self-assemble into bacterial microcompartments (BMCs) that encase the enzymes necessary for EA metabolism. For the first time, native EUT BMCs were fluorescently tagged, and EUT BMC formation was observed in vitro, and in vivo. Interestingly, BMC formation was observed in bacteria infecting Caco-2 cells, but not the macrophage cell lines. Finally, the cellular immune response of Caco-2 cells to infection with *eut* mutants was examined, and it was discovered that *ΔeutB* and *ΔeutV* mutants similarly elevated the expression of inflammatory cytokines. In conclusion, EA sensing and utilization during LMO intracellular infection are important for optimal LMO replication and immune evasion but are not always concomitant with BMC formation.

## INTRODUCTION

Ethanolamine (EA) is a metabolically useful compound that arises from the membrane lipid phosphatidylethanolamine contained in all bacterial and eukaryotic cells (1, 2). Because of the natural turnover of the epithelium and the cells of the microbiota, the gastrointestinal environment is particularly rich in EA with concentrations as high as 2mM (3, 4). As this compound can serve as a useful source of carbon and/or nitrogen, there is a correlation between bacteria that contain the genes to catabolize this EA and those found in the gut including species of *Enterococcus, Escherichia, Clostridium, Salmonella* and *Listeria* (reviewed by (5, 6)). However, expression of the *eut* genes has been documented in other host environments, including the intracellular one, and may be of relevance for *Listeria monocytogenes* (LMO) (7, 8).

LMO is a facultative, intracellular pathogen, capable of invading, surviving, and multiplying inside various host cell types including cells of the gastrointestinal epithelium and the innate immune cells. Upon attachment, invasion is mediated by engulfment and internalization into a vacuole. Depending on the host cell environment, LMO can disrupt the phagosome, escape into the host cytoplasm, replicate, and spread using actin-based motility (reviewed by (9)). In the host environment of human colon epithelial (Caco-2) cells, a microarray analysis found that the *eut* operon of LMO was upregulated (8). Furthermore, a study of LMO gene expression under several different host conditions, discovered strong upregulation of the *eut* genes in the intestines of infected mice and weaker, but still significant levels of expression in human blood (7).

The EA utilization genes are usually found together in an operon or locus and encode the necessary regulators, enzymes, and structural proteins (5, 6)). Specifically, *Listeria monocytogenes* possesses 17 genes involved in EA metabolism (Fig. 1A) (10). The regulation of the *eut* genes for this bacterium and other Gram-positives such as *Enterococcus* involves an EA-responsive sensor kinase, encoded by *eutW,* a response regulator of the ANTAR family, encoded by *eutV,* and an AdoCbl-dependent riboswitch (10–13). To metabolize EA, an ammonia lyase, composed of two sub-units encoded by *eutBC*, is required for the initial catabolic step that generates acetaldehyde and ammonia. To concentrate these products for the downstream reactions and to prevent interaction of toxic acetaldehyde with the cytoplasm, some, including LMO, also encode for structural proteins that self-assemble into enclosed, proteinaceous structures called bacterial microcompartments (BMCs) (reviewed by (5, 6)).

**Fig. 1:**
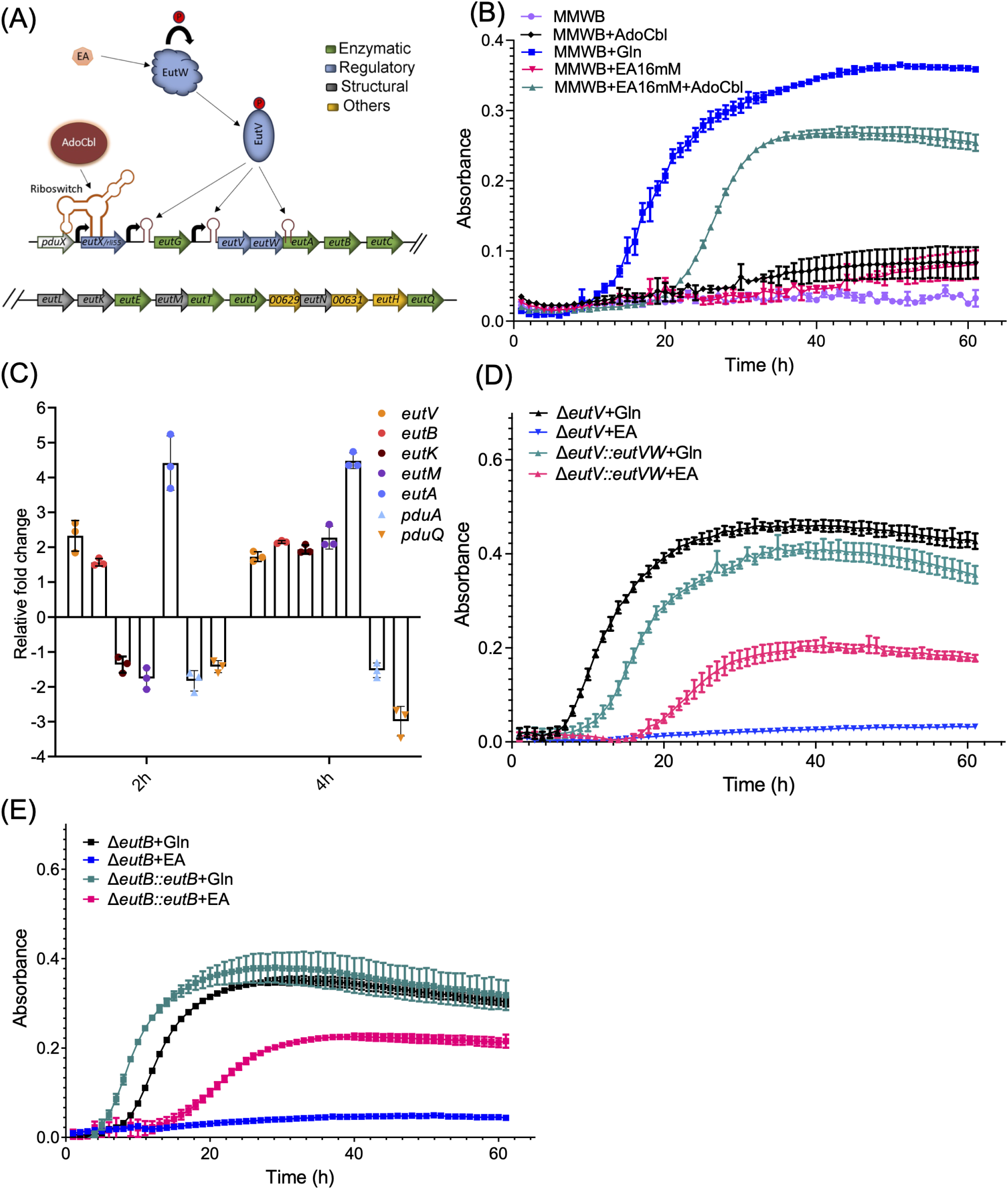
MMWB plus EA supports LMO10403s growth dependent on *eutV* and *eutB.* (A) Graphical representation of the organization and regulation of the *eut* operon in LMO. See text for details. (B) Growth curve of LMO when grown in MMWB containing either glutamine (Gln) or ethanolamine (EA) as a source of nitrogen. (C) Relative expression of the *eut* genes when LMO bacteria were grown in the presence of EA with respect to Gln. (D) Growth curve comparison of *ΔeutV and ΔeutV* complemented *(ΔeutV::eutVW)* when grown in the presence of Gln or EA. (E) Growth curve comparison of *ΔeutB and ΔeutB* complemented *(ΔeutB::eutB)* when grown in the presence of Gln or EA. Shown is the average of three replicates and error bars represent the standard deviation (SD).

In addition to upregulation of the *eut* genes, a contribution of ethanolamine metabolism to LMO intracellular replication in Caco-2 cells was noted; a *eutB* mutant was deficient compared to the parental strain (8). Furthermore, in an intravenous (i.v.) mouse infection model, a *eutB* mutant was deficient in dissemination to the spleen and liver (10). Since EutB is one of the subunits that comprise the ammonia lyase, these data suggest EA catabolism promotes survival in both gastrointestinal epithelial cells and during bloodstream infection (8, 10). Interestingly and in contrast, replication of a *eutB* mutant strain of *Salmonella enterica* serovar Typhimurium was not deficient in host cells. However, replication was diminished for a strain deleted for *eutR*, which encodes the EA-sensing transcriptional regulator (14). This was attributed to the fact that EutR regulates genes associated with pathogenicity in *S.* Typhymurium – the SPI-2 pathogenicity island. Further work by the Kendall group established the importance of the EutH permease for *S.* Typhimurium (15). EutH is required for EA entry under acidic conditions such as those encountered in the phagosome (16). Interestingly, a slight phenotype for a *eutH* mutant in intracellular replication was also shown for LMO, but regulatory or metabolism mutants were not directly compared (15). Whether a LMO *eutV* mutant strain would display a greater deficiency than a *eutB* mutant strain in intracellular replication like *S.* Typhimurium is unknown.

As mentioned above, EA metabolism is frequently, but not always encased in an organelle-like structure called a bacterial microcompartment (BMC) (reviewed by (6, 17, 18)). Rather than having a selectively permeable membrane, these “organelles” have a selectively permeable protein shell made up of structural components. A feature of the assembly is that some of the enzymatic proteins have “encapsulation peptides” (EPs) that interact with the structural proteins to enable BMC formation. Native EUT BMCs have been visualized in several species of bacteria including LMO by transmission electron microscopy (TEM) (19). However, fluorescent-tagging of EUT BMCs to our knowledge was only done in a heterologous system in which *eut* genes encoding the structural proteins of *S.* Typhimurium were over-expressed in *E. coli* using a constitutively active promoter, and BMCs were visualized by tagging an encapsulation peptide with GFP (20–22). Here we report the fluorescent tagging of native EUT proteins under the control of *eut* promoters to permit the visualization of endogenous BMC formation in vitro and in host cells.

In this work, we sought to further characterize the role of EA utilization during LMO infection of host cells. Unlike what was previously observed with *S.* Typhimurium (14), an EA metabolism mutant (*ΔeutB*) was equally deficient in intracellular replication compared to an EA regulator mutant (*ΔeutV),* and this was consistent across a number of cell types. To visualize EUT BMC formation, native structural proteins were fluorescently tagged and expressed under *eut* promoters. EUT BMC formation was observed in vitro, and for the first time, in vivo, in some, but not all host cells tested. Interestingly, formation of EUT BMCs was associated with the phagosomal phase of infection. Finally, it was found that LMO *ΔeutB* and *ΔeutV* mutants similarly affected the cellular immune response; comparable increases in several inflammatory cytokines were observed.

## RESULTS

### Establishment of conditions in which LMO can utilize ethanolamine (EA) as a nitrogen source dependent on the *eut* genes

Previous work demonstrated that LMO can utilize EA as a source of carbon and/or nitrogen depending on the media conditions (19, 23). We optimized a similar chemically defined medium that we named MMWB (Modified MWB) (Table S1) (24–27). MMWB contained 4.1mM glutamine (Gln) as the nitrogen source, allowing for robust growth of LMO (Fig. 1B). EA could be substituted for Gln, which requires the bacteria to utilize EA as a nitrogen source. As expected, no growth was observed in the absence of Gln, underscoring the necessity of a nitrogen source (Fig. 1B). To test if EA could substitute as a nitrogen source in the medium, we replaced Gln with 16mM EA. However, no growth was observed until we additionally supplemented the medium with 200µM of AdoCbl (Fig. 1B). The data aligns with previous research demonstrating a requirement for AdoCbl as a co-factor in the metabolism of EA and in the induction of the *eut* genes (reviewed by (6)), and all subsequent experiments with media conditions labeled “EA” included AdoCbl. In conclusion, we show that LMO can grow in MMWB utilizing EA as a nitrogen source.

As described in the introduction and illustrated in Fig. 1A, the *eut* locus of LMO contains 17 genes that encode the regulators, enzymes, and structural shell proteins necessary for EA metabolism. To verify induction of these genes in MMWB + EA, expression of a subset was measured by qRT-PCR. As expected, all genes measured, *eutV, A, B, K,* and *M,* displayed increased expression following 4 hours of growth in MMWB + EA relative to MMWB + Gln. Of note, the expression was specific for *eut* genes associated with EA metabolism; genes associated with propanediol (PDU) metabolism, *pduA* and *pduQ,* were not induced (Fig 1C).

To further verify the role of the *eut* genes in LMO’s use of EA as a nitrogen source in MMWB, we generated strains containing *ΔeutV* (LMAC030) and *ΔeutB* (LMAC031) non-polar, in-frame, deletion mutations in the parental background (Fig 1D and 1E). When grown in the presence of Gln as the nitrogen source, these mutant strains exhibited growth comparable to the parental strain. However, they did not grow when EA was provided as a nitrogen source, indicating that the response regulator, EutV, and the ammonia lyase subunit, EutB, were indispensable for growth in the presence of EA in vitro, as expected. To ensure that the defects in growth were due to the introduced mutations, we complemented these strains. In order to retain EA-inducibility, the complementation constructs included the regulatory sequences upstream of *eutV* or *eutG.* Both these regions contain predicted promoters followed by ANTAR binding sites that are the hallmarks of antitermination regulation by EutV (Fig 1A) (10). Growth in EA was partially restored for the *ΔeutV* strain when *eutV,* under the control of the *eutV* promoter, was introduced as a transgene into a chromosomal site outside of the EA locus (see Materials and Methods for details), (Fig 1D and S1A). Note that successful complementation of *ΔeutV* required using the combined *eutVW* sequence; attempts to complement with the *eutV* sequence alone were unsuccessful. As the *eutV* and *eutW* genes overlap and appear to be co-transcribed, it is possible that a *eutV* transcript alone is unstable. Growth in EA was also restored in the *ΔeutV* background, but to a lesser degree, when the *eutG* promoter was utilized (Fig S1A). In contrast, successful complementation was only achieved in the *ΔeutB* strain with the *eutG* promoter driving the expression of *eutB*; use of the *eutV* promoter did not enable growth in EA (Fig 1E and S1B). We speculate that these differences in the promoters’ abilities to restore growth in EA is due to differences in the levels of EutV and EutB that are required for EA metabolism and achieved by the promoters. Overall, these results established conditions in which LMO can utilize EA as a source of nitrogen dependent on the positive regulation of the *eut* genes (EutV) and the ability to metabolize EA (EutB).

### LMO EUT BMC formation is dependent on EA, EutV, and EutB

EA utilization has been associated with BMC formation for many EA-utilizing bacteria including *L. monocytogenes* (6, 19). Specifically, LMO EUT BMCs were previously visualized under anaerobic conditions (19). To test for BMC formation in aerobic MMWB + EA conditions, transmission electron microscopy (TEM) was utilized. Structures resembling BMCs were readily visible in the parental LMO background following 6 hours growth in medium containing EA but not Gln (Fig. 2A) with 35.3% of the cells examined (n=51) containing at least one visible BMC. (Note that this count is likely an underestimate due to the fact that a given plane of BMC-containing cell may not contain a BMC.) As dependence of the formation of these structures on EutV and EutB had not been previously reported in LMO, we also examined *ΔeutV* and *ΔeutB* deletion mutant strains grown in EA; both strains failed to form any visible BMCs (Fig. 2B and 2C). The complemented strains both restored BMC formation, as predicted, indicating that the deletions did not have polar effects on the expression of the downstream structural genes (Fig. 2B, 2C and 1A). Specifically, 37.5% and 38.4% of the *eutV* (n=24) and *eutB* (n=26) complemented cells presented with at least one visible BMC.

**Fig 2:**
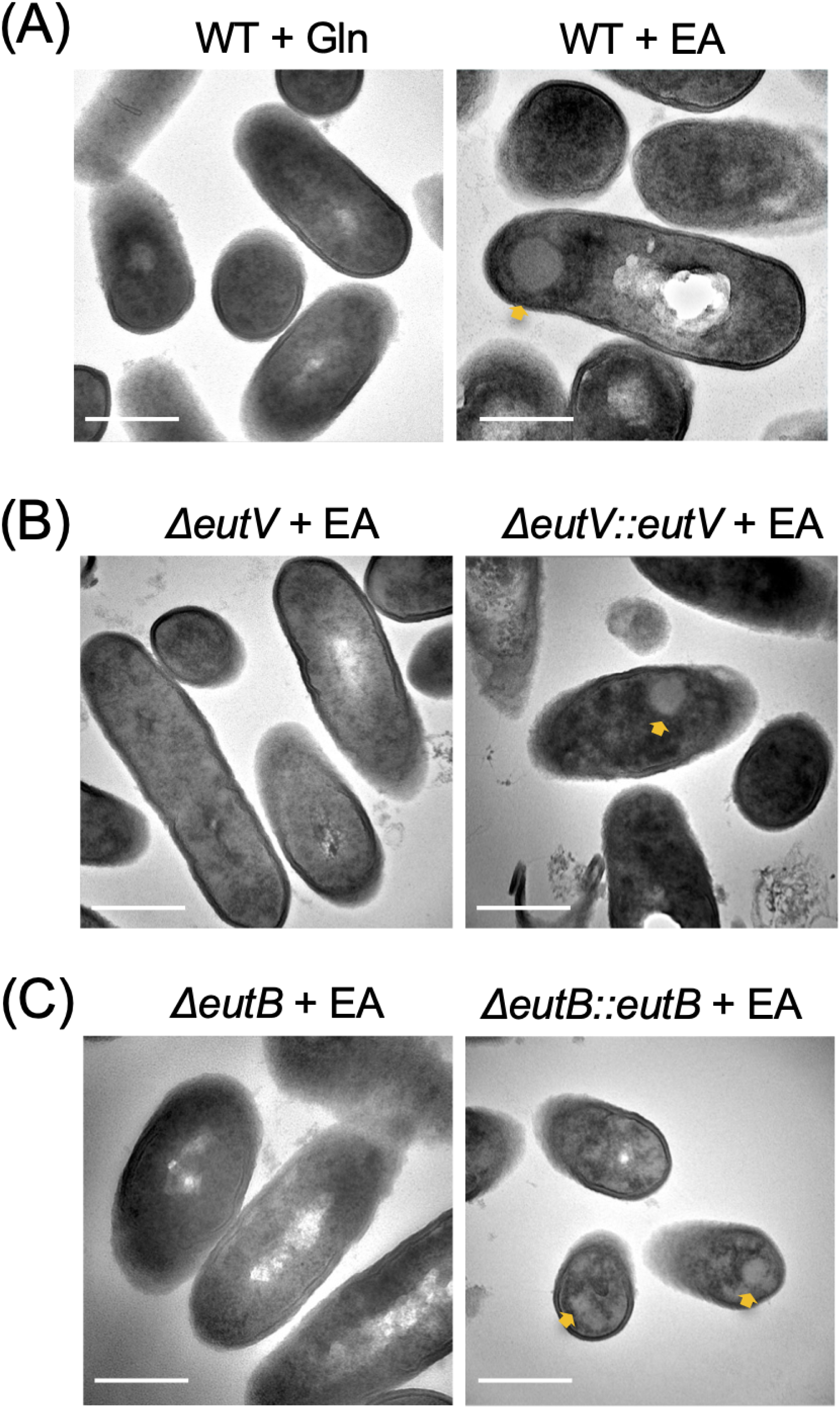
EUT BMCs form in medium containing EA. TEM images showing the formation of EUT BMCs when LMO was grown in MMWB with (A) Gln or EA. No such structures were visible in the *ΔeutV* (B) and *ΔeutB* (C) strains grown under the same conditions. Complementation of *ΔeutV* and *ΔeutB* restored the formation of BMC structures (B and C right panel respectively). Scale bar represents 500nm and yellow asterisks indicate the BMCs.

In order to visualize EUT BMC formation in live cells, we introduced a *eutK::mCherry* translational fusion under the control of the EA inducible *eutG* promoter into an ectopic, chromosomal site in wild-type, *ΔeutV* and *ΔeutB* backgrounds (Materials and Methods). Tags were not added to the endogenous structural genes, as it was previously reported that this can disrupt BMC formation (28, 29). To ensure that introduction of the transgene did not disrupt the ability of the strain to utilize EA, growth curves were performed, and the strain was found to grow in EA with kinetics similar to the parental strain (Fig. S1C). Following six hours of incubation in medium containing EA, distinct red punctate structures were observed corresponding to BMC-like formations in about 85% of the cells (Fig. 3A and 3E). Notably, when bacterial cells were cultivated in a medium containing Gln, these BMC-like structures were rarely evident under confocal microscopy. (Fig. 3B and 3E). By performing live cell imaging of this strain, the dynamics of these structures forming and dissipating was revealed (Mov. S1). Consistent with the TEM observations, BMCs were not visible in the *ΔeutV* and *ΔeutB* strain backgrounds (Fig. 3C, 3D and 3E). In addition to the *eutK::mCherry* translational fusion, *eutM::mCherry* and a *eutK::mNeonGreen* fusions were additionally generated. They presented with a similar punctate fluorescent pattern when grown in EA, indicating that the observed localization was unlikely to be artifactual due to a specific fusion or tag (S2A and S2B). In conclusion, these experiments demonstrate that EUT BMC formation is dependent on EA, EutV, and EutB and can be visualized with fluorescent tags in *L. monocytogenes*.

**Fig 3:**
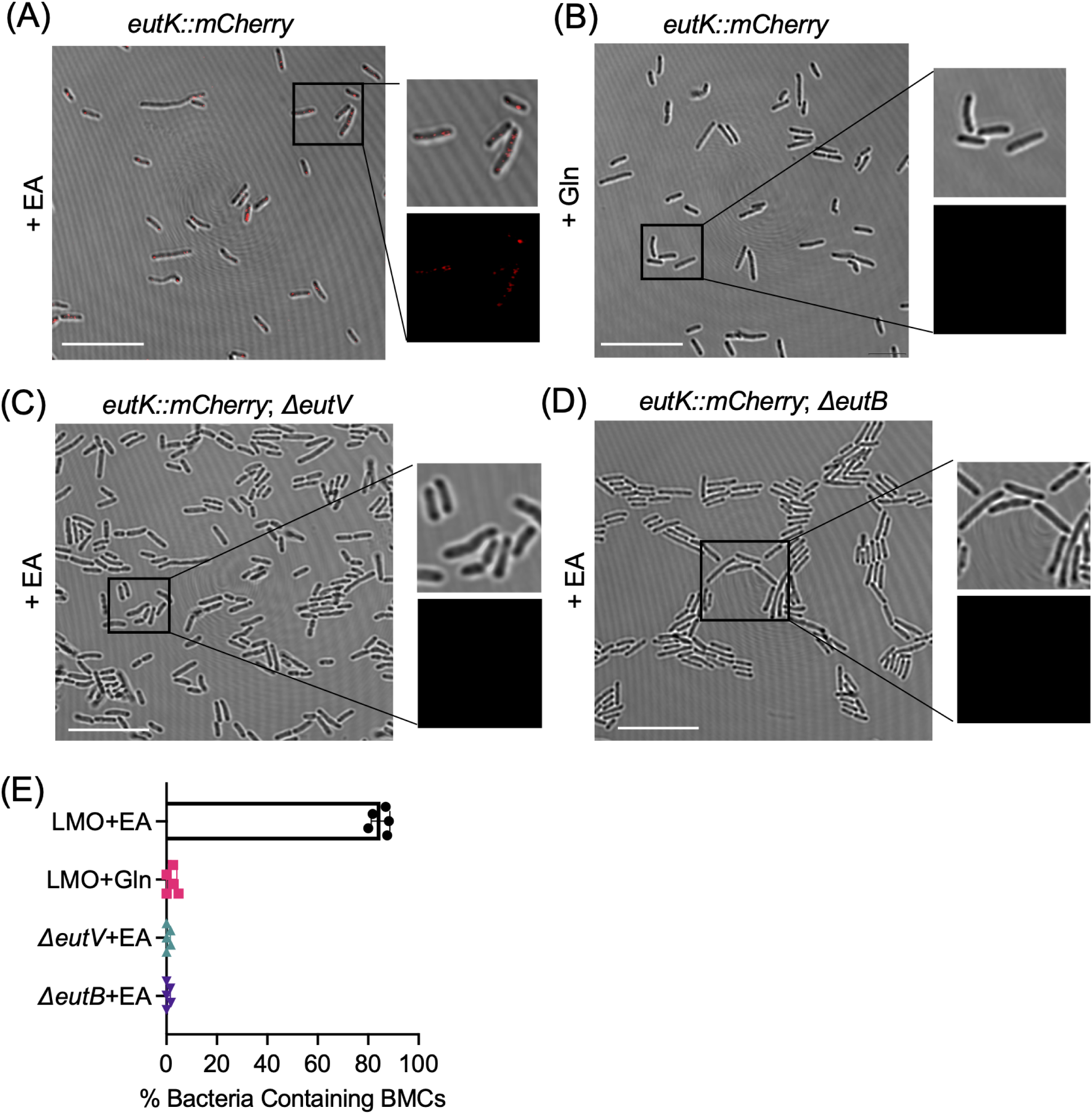
EUT BMCs can be fluorescently tagged. Confocal images of the LMO *eutK::mCherry* strain when grown in in MMWB with either EA (A) or Gln (B) as a nitrogen source. (B) Confocal images of LMO strains containing the *eutK::mCherry* transgene (red) in *ΔeutV* (C) and *ΔeutB* (D) backgrounds following growth in MMWB with EA. Scale bar represents 10µm. (E) Percentage of cells with visible BMCs. Shown is the average from replicates containing an n of 20-60 cells and the error bars represent the SD.

### Loss of EA utilization impairs intracellular replication of LMO

As LMO invades the cells of the gastrointestinal epithelium in humans, the Caco-2 human colorectal adenocarcinoma cell line is frequently used to model LMO intracellular infections (30, 31). Previous work on strain LMO EGDe documented that a strain with an insertion mutant in *eutB* was deficient in Caco-2 intracellular replication (8). To verify the phenotype with LMO 10403S, we infected Caco-2 cells with the parental strain and the *ΔeutV* and *ΔeutB* strains. Following a one-hour infection at an MOI of 5, the cells were washed, and medium containing gentamycin was added for one hour to kill extracellular bacteria. At the zero-time point, defined as when the extracellular bacteria were removed and the cells washed, the number of intracellular bacteria between the strains was not significantly different, indicating no defect in bacterial invasion (Fig. 4A). However, by three hours, there was approximately half-a-log less CFUs of the *ΔeutV* and *ΔeutB* strains compared to the parent, but no significant difference in the level of attenuation between the two mutant strains. Note, that at this time point there is a drop in the number of bacteria compared to the zero-time point. It is thought that CFUs at the zero-time point can be inflated due to bacterial adherence to the surface of Caco-2 cells despite the wash step, and this phenomenon has been observed previously during LMO infection (30). By six hours, the CFUs were increased over the zero-time point, indicative of intracellular replication, but the attenuation of the *eut* mutants persisted (Fig. 4A).

**Fig. 4:**
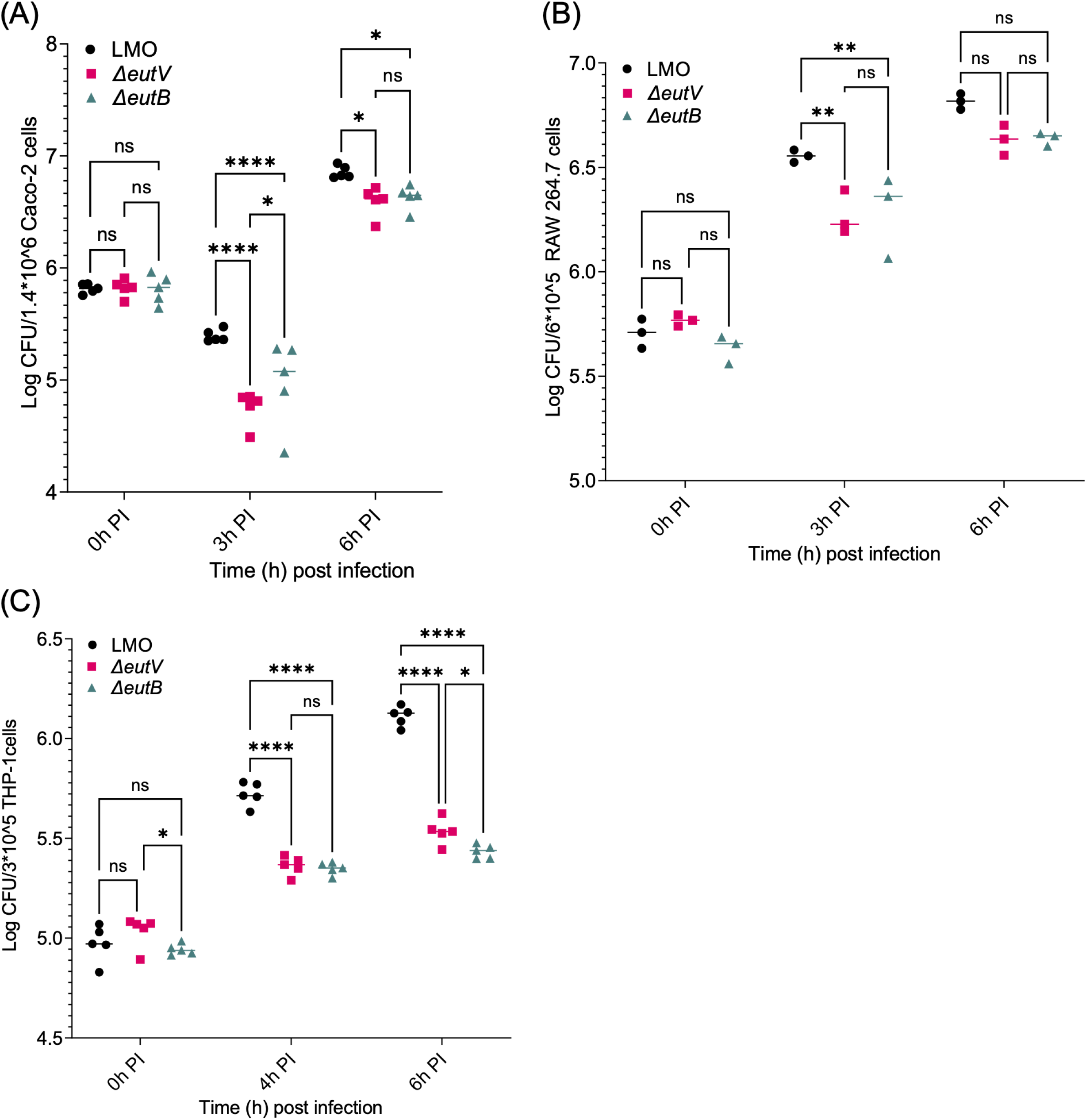
*eut* mutants are deficient in intracellular replication: Bacterial CFUs were quantified in Caco-2 cells (A), RAW264.7 macrophages (B) and M1 polarized THP-1 macrophages (C). At the indicated time points post-infection, the cells were washed, lysed, and serial dilutions of the lysates were plated as described in methods. The mean of three replicates were calculated with the error bars indicating the standard deviation. 2-way ANOVA with Bonferroni multiple testing correction was used to calculate the significance of the indicated comparisons. *P < 0.05; **P < 0.01; ***P < 0.001; ****P < 0.0001.

Following enterocyte invasion, *L. monocytogenes* can infect recruited immune cells such as macrophages, which can lead to systemic infections (9). A LMO mutant in *eutH,* which encodes for an EA transporter thought to be necessary for EA utilization in acidic environments, was previously shown to have a slight defect in intracellular replication in bone marrow-derived macrophages (BMDMs), suggesting that EA utilization might play a role in this process (15). To further test the efficiency of intracellular replication of the *eut* mutant strains in immune cells, we infected RAW 264.7 and differentiated THP-1 cells, which are mouse and human derived macrophages, respectively (Fig. 4B and 4C). Replication of the *ΔeutV* and *ΔeutB* strains was attenuated by about 0.5 logs following three-four hours of infection, and the attenuation persisted at the six-hour time point. Again, there was no significant difference in the degree of deficiency between the two strains. In conclusion, the *ΔeutV* and *ΔeutB* strains are deficient in intracellular replication to a similar degree in a range of cell types suggesting that EA utilization positively contributes to LMO infection efficiency.

### LMO forms EUT BMCs during intracellular infection of Caco-2 cells

To test for BMC formation in the intracellular environment, Caco-2 cells were infected with the strains expressing the *eutK::mCherry* fusion. Strains containing the *eutK::mCherry* translational fusion were utilized for the in vivo experiments because they were brighter compared to the *eutM::mCherry* strains (Fig. S2A) and were less prone than mNeonGreen to interference from background autofluorescence (Fig. S2B). The samples were fixed at 2, 4, 6 and 8 hours post-infection, and bacterial cells were visualized using an antibody specific to *L. monocytogenes.* Host cell actin was visualized by staining with phalloidin. At 4, 6, and 8 hours, punctate, red fluorescence inside some of the bacterial cells from the parental strain was visible, indicative of BMC formation (Fig. 5A and Fig. S3A). As expected, little to no signal was observed in samples from the *ΔeutV* and *ΔeutB* strains containing the *eutK::mCherry* fusion (Fig. 5B-C and Fig. S3A). Time points later than 8 hours were not captured due to host cell lysis. However, as shown by imaging the larger fields (Fig. S3A), not all bacteria within the host cells presented with visible BMCs. By counting the number of bacteria presenting with and without BMCs in a given image, it was determined that approximately 25% of the *eutK::mCherry* parental strain of LMO had BMCs. However, the percentage of bacteria containing BMCs in a given image was variable and became more so over time (Fig. S3B). As host cell actin will associate with cytoplasmic LMO, escape from the phagosome can be visualized by the formation of actin clouds and tails (Fig. 5D). Interestingly, we observed that bacteria associated with actin clouds or tails never contained visible BMCs. Rather BMC formation was observed in some, but not all, bacteria unassociated with actin (Figure 5D and S3A). Note that the signal from the antibody against LMO (green) was weaker in the actin associated bacteria, perhaps due to actin interfering with antibody binding. Therefore, we have outlined some of the bacteria with actin tails in Fig. 5D to aid in visualization. Actin tails and clouds also formed during infection with both the *eut* mutant strains, demonstrating that phagosomal escape is not blocked (Fig S3A). In conclusion, these data show that in Caco-2 cells EUT BMCs form in LMO. Moreover, because BMCs were only observed in some bacteria not associated with actin, the results suggest that BMC formation may occur preferentially in the phagosome.

**Fig. 5:**
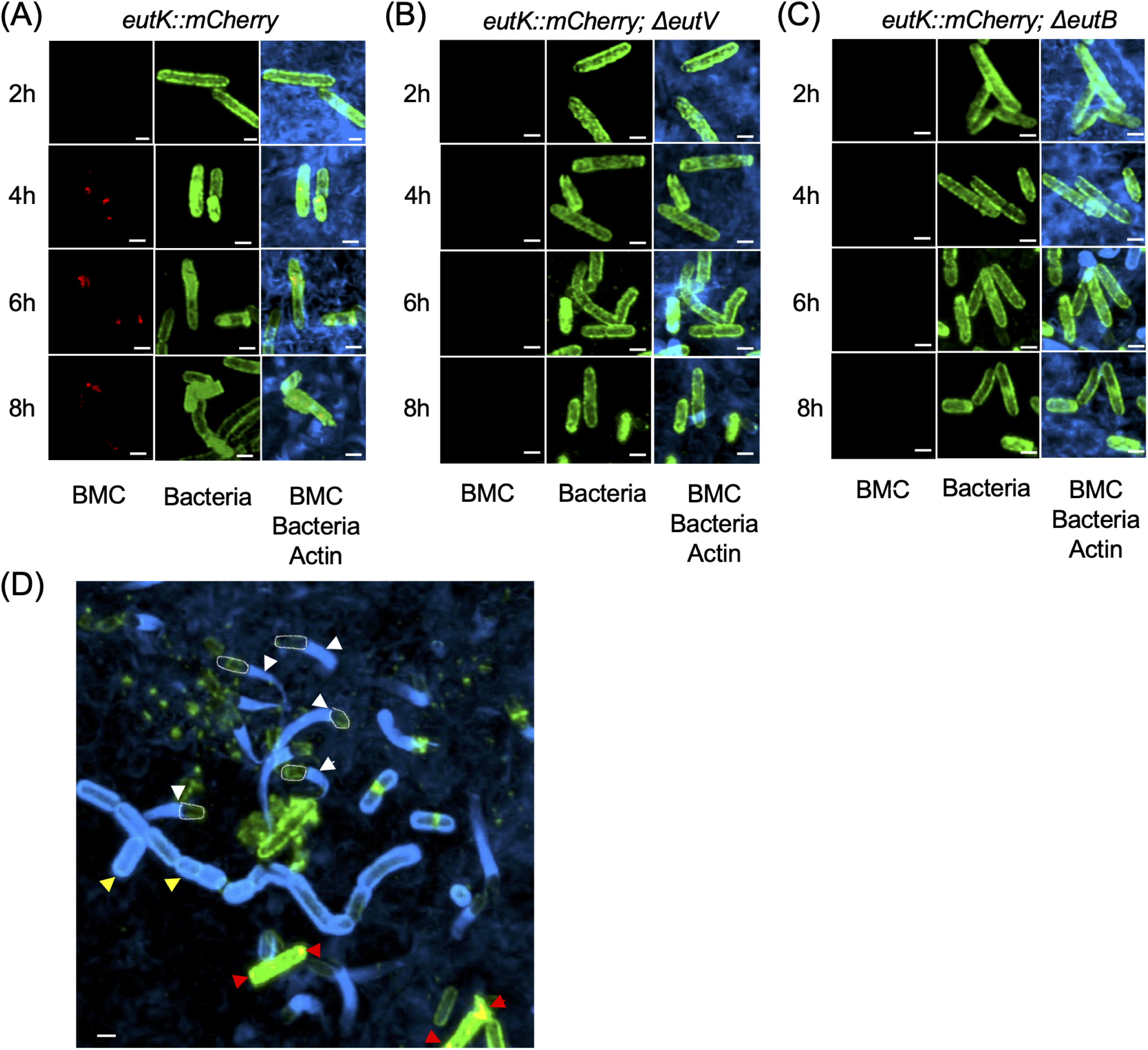
LMO forms visible BMCs in Caco-2 cells dependent on *eutV* and *eutB*. Caco-2 cells were infected with LMO strains containing the *eutK::mCherry* transgene (red) in the wild type (A), *ΔeutV* (B) and *ΔeutB* (C) backgrounds. The cells were fixed at the indicated time points and stained with phalloidin to visualize actin (blue) and α-LMO antibody to visualize the bacteria (green). Representative confocal images are shown. Scale bars represents 1µm. (D) Close-up of boxed view of LMO containing *eutK::mCherry* at the 6 hour time point from Fig. S3. Red arrows indicate BMCs. Yellow arrows indicate examples of bacteria surrounded by an actin cloud. White arrows point to examples of actin tails. Five bacteria associated with an actin tail have been outlined in white for easier visualization.

Confocal imaging was also performed on RAW 264.7 cells infected with the *eutK::mCherry* strain at 2, 4, and 6 hours post-infection (Fig. S4A). Following staining and imaging as described for the Caco-2 cells, actin tails and clouds were observed at later time points, typical of intracellular LMO infection. However, despite the *eut* genes contributing to intracellular replication in these cells (Fig. 3B), we were unable to detect any mCherry signal at any time point (Fig. S4A). The same experiment was performed using M1 differentiated THP-1 cells, reported to delay phagosomal escape (32). No mCherry signal was evident during the infection of these cells at either early or late time points (Fig. S4B). However, normal infection kinetics were observed with actin clouds and tails visible by five hours. These results suggest that BMCs do not form in RAW 264.7 or THP-1 cells.

### Loss of EA utilization increases host cell inflammatory responses

To further investigate the role of EA utilization in LMO’s interaction with host cells, we examined the inflammatory response of Caco-2 cells following infection. Specifically, the expression of genes encoding for inflammatory cytokines, vital signaling mediators that orchestrate and amplify the immune response by recruiting immune cells and enhancing antimicrobial capabilities, were measured (33, 34). Caco-2 cell cultures were subjected to infection with the LMO parental or the *ΔeutV* and *ΔeutB* mutants as before, and host cell RNA was extracted at four hours post-infection. The expression levels of specific inflammatory cytokines, interleukin-1 beta (IL-1β), tumor necrosis factor-alpha (TNF-α), interleukin-6 (IL-6), interleukin-8 (IL-8), and chemokines CXCL2 and MCP-1 were examined by qRT-PCR as described previously for Caco-2 cells (35). A 2-4-fold increase in the expression of these factors by Caco-2 cells infected with the *ΔeutV* and *ΔeutB* mutant strains was observed in comparison to those infected with the parental strain (Figure 6). There were no significant differences in the expression levels of these cytokines between the *ΔeutV* and *ΔeutB* mutant strains, suggesting that the observed differences are solely due to the loss of ethanolamine utilization. The stronger inflammatory response coupled with the reduced intracellular replication observed during infection with the mutant strains (Figure 3A) suggests that loss of ethanolamine utilization reduces LMO’s ability to evade the host immune system.

**Fig 6:**
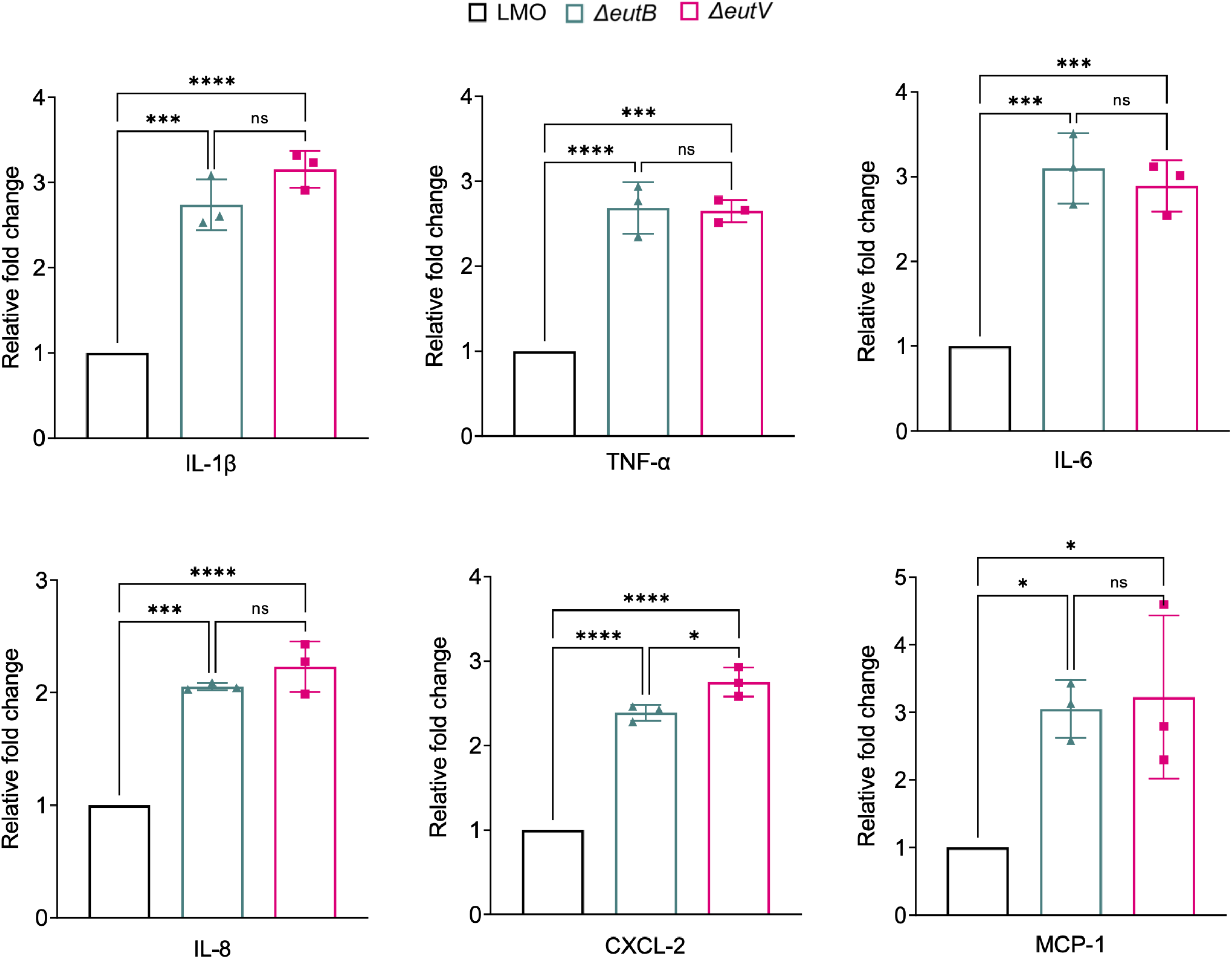
*eut* mutants increase pro-inflammatory signatures during Caco-2 infection. Following infection of Caco-2 cells with the indicated strains, host cell RNA was isolated and qRT-PCR analysis revealed a significant elevation in the expression of genes encoding inflammatory cytokines interleukin-1 beta (IL-1β), tumor necrosis factor-alpha (TNF-α), interleukin-6 (IL-6), interleukin-8 (IL-8), and chemokines CXCL-2 and MCP-1. The mean of three replicates were calculated with the error bars indicating the standard deviation. 2-way ANOVA with Bonferroni multiple testing correction was used to calculate the significance of the indicated comparisons. *P < 0.05; **P < 0.01; ***P < 0.001; ****P < 0.0001.

## DISCUSSION

In this study, we demonstrated that the catabolism of EA contributes to the efficacy of intracellular infection of LMO in a variety of host cells. In *S.* Typhimurium, by contrast, the ability to sense, but not catabolize EA, was shown to enhance intracellular survival in macrophages; a *eutR* regulatory mutant was deficient in intracellular survival, but a *eutB* metabolic mutant was *not.* (14). Here, a direct comparison between LMO *eutV* regulatory and *eutB* metabolic mutants revealed indistinguishable phenotypes. Both exhibited the same level of depressed intracellular replication and caused similar changes to the host cell immune response (Fig. 3 and 6). The difference could be related to the scope of the genes regulated by EutR and EutV. EutR is a DNA binding transcriptional regulator. In addition to binding its promoter in the *eut* locus, EutR directly binds the promoter of the *ssrB* gene in *S.* Typhimurium encoding a transcriptional regulator. SsrB activates the expression of the Salmonella pathogenicity island 2 (SPI-2) genes that are required for *S*. Typhimurium intracellular replication and survival (14). EutR in EHEC also regulates virulence related genes outside of the *eut* locus (36, 37). In contrast, EutV is an ANTAR (AmiR and NasR transcriptional antiterminator regulators) response regulator that binds specific sequences in mRNA leaders that overlap terminators, preventing their formation (38, 39) In LMO, there is no evidence that EutV directly regulates genes other than those related to EA utilization (10, 38). The observation that strains containing *eutV* and *eutB* deletions caused identical effects on LMO intracellular replication as well as on host cell immune responses is consistent with the phenotypes being caused by the loss of the ability to catabolize EA in this bacterium, unlike in *S.* Typhimurium. How exactly the loss of EA metabolism causes these changes in LMO’s interactions with host cells is of future interest.

By fluorescently tagging BMC structural proteins, we were able to visualize BMC formation in vitro and in vivo. We did observe BMCs forming in a subset of the bacteria inside Caco-2 cells. Interestingly, these visible BMCs were only in seen in bacteria not associated with actin; there were no examples of BMCs in bacteria associated with tails or clouds in our imaging experiments. These data suggest that BMCs might only form during the phagosomal phase of infection. However, attempts to confirm this observation with different approaches such as antibodies against phagosomal markers and/or developing antibodies to visualize the BMCs by immunolabeling, were unsuccessful.

Unlike the Caco-2 cells, no BMCs were observed during LMO infection of RAW 264.7 and THP-1 cells. We postulate two possible explanations for the lack of visible BMC formation in these monocyte/macrophage cell lines. One, BMCs were generated, but were unable to be visualized due to low expression of the *eutK::mCherry* transgene. Two, the metabolism of ethanolamine is occurring at a lower level in these cells that does not require BMC formation. Recall that some bacterial species, like *Pseudomonas aeruginosa* and *Acinetobacter baumannii* encode for the core EA metabolic enzymes, but lack the structural proteins (39, 40). Even in bacteria such as *S. typhimurium* that contain the full complement of structural proteins, it has been shown that they are still able to metabolize EA, albeit with less efficiency, when they are deleted for the microcompartment-encoding genes (41, 42). Perhaps there are differences in the amount of free EA available in these different host cells that determine whether BMC formation is necessary for EA utilization or not. Future work using transcriptomics and metabolomics to further understand the host cell environments and the bacterial response may provide more insight into these observations.

Another surprising finding in this study was the lack of BMC formation in the *ΔeutB* mutant. We expected that no structures would form in the *ΔeutV* mutant that lacks the regulator necessary for expression of all the *eut* genes but were surprised that a strain missing a single catalytic enzyme failed to form BMCs. However, it has been reported that the shell proteins of the carbon-fixing β-carboxysomes condense around key catalytic enzymes necessary for nucleation of the BMC structure (18, 29). In contrast, the α-carboxysomes can assemble either shell-first or undergo concomitant shell–core assembly, and the propanediol (PDU) metabolizing BMCs form independent cargo and shell aggregates (43–45). While there have been studies in heterologous systems to examine the requirements for EUT BMC assembly (20–22), the dynamics have not been determined in a natural system. Therefore, it is possible that core assembly requiring EutB is necessary for shell protein condensation. However, our previous work in *E. faecalis* documented normal BMC formation by TEM in strains with *eutB* mutations (46), raising the possibility of species-specific differences. Undoubtedly, it will be of interest to further study EUT BMC formation and elucidate the assembly pathway(s). More broadly, the field should investigate if the assembly of BMCs varies in a manner dependent on species and environmental context, including host environments. In conclusion, EA utilization contributes importantly to LMO intracellular replication, but may not always require EUT BMC assembly.

## MATERIALS AND METHODS

### Bacterial strains and media

*Listeria monocytogenes* 10403s (LMO) strain (source: Bei resources) was used in this study and all mutations and recombinants were constructed in this background. All LMO strains were grown in brain heart infusion (BHI) broth or on BHI agar (1.5% agar) plates supplemented with streptomycin (50 µg /ml) in 37 °C. The chemically defined medium MMWB (Table S1) was prepared by modifying the original recipe reported by Welshimer (24). Medium was always prepared fresh, and filter sterilized with a 0.22micron filter. MMWB medium was supplemented with either glutamine (Gln: 4.1mM) or ethanolamine (EA: 16mM) and VitB12 (200µM) as needed. Cloning was done in either *E. coli* Top10 or DH5α. All *E. coli* strains were grown in Luria-Bertani (LB) broth and were supplemented with antibiotics [Ampicillin (100 µg /ml) or Kanamycin (50 µg/ml)] wherever required. For recombinant LMO strains, appropriate antibiotics [Kanamycin (50 µg/ml) and streptomycin (50 µg/ml)] were used.

Generation of recombinant, *ΔeutV*, *ΔeutB* and the various complemented strains in LMO All primers used in this study were purchased from Sigma-Aldrich and are listed in Table S2. PCR reactions were performed using Platinum™ SuperFi II DNA Polymerase (ThermoFisher Scientific: 12361010) according to the manufacturer’s protocol. Restriction free cloning was conducted using NEBuilder® HiFi DNA Assembly (NEB: E2621L). Isolation of plasmids and purification of PCR products were conducted by using commercially available kits from either Qiagen or Macherey-Nagel. Sequencing was conducted at Genewiz, and the results visualized and verified using SnapGene (version 5.2.5.1). For creating the *ΔeutV* (LMAC030) and *ΔeutB* (LMAC031), the respective upstream and downstream regions were cloned into pMAD and electroporated into LMO as reporter earlier (47). Electrocompetent cells of LMO were prepared as described (48). The complemented strains *ΔeutV::eutV* (LMAC037) *and ΔeutB::eutB* (LMAC034) were created by cloning the respective gene along with promoter region of either *eutV* or *eutG* in pIMK plasmid followed by electroporation. For creating the recombinant strain expressing mCherry tagged to EutK (LMAC038), the *eutK* coding sequence was driven by the *eutG* promoter and fused to the mCherry coding sequence. It was then cloned into the pIMK plasmid to create the construct AC038 followed by electroporation as described by Monk *et al*, 2008. Similarly, AC038 was electroporated into LMAC030 (*ΔeutV*) and LMAC031 (*ΔeutB*) to generate strains LMAC040 (*eutK::mCherry*; *ΔeutV*) and LMAC043 (*eutK::mCherry*; *ΔeutB*), respectively.

### Growth curves

LMO and the mutants were grown overnight in 10 ml of BHI broth at 37°C with aeration. Bacterial cells were harvested and washed thrice with PBS followed by a wash with MMWB medium devoid of glutamine or EA. Cultures were then seeded on a 48 well plate from Falcon® (Corning) containing the desired medium with a starting OD_600_ of 0.05. Plates were sealed with Breathe-Easy® sealing membranes (Diversified Biotech) and the growth curve was monitored on a Cytation™ 5 plate reader system (BioTek Instruments Inc) for 72 hours, with OD readings taken every hour.

### Transmission Electron Microscopy

Bacterial strains were grown in BHI broth to midlog phase at 37°C with aeration. Bacterial cells were then harvested, washed, and processed as mentioned above. The strains were resuspended in MMWB medium containing either Gln (4.1mM) or EA (16mM) and Vit B12 (200µM) as needed. Cells were then harvested, fixed, processed and sliced (12). Imaging was conducted in JEOL 1400 TEM system, equipped with the Gatan 2k x 2k CCD camera, and accompanying software.

### Cell culture and infection

All the cell lines were purchased from Sigma. The murine macrophage RAW264.7 cell line and Human colon adenocarcinoma Caco-2 cell line was cultured in DMEM (Gibco: 11995065) supplemented with 10% fetal bovine serum (FBS) (Gibco: 10082147) at 37°C in a humidified atmosphere at 5% CO_2_. For culturing the Caco-2 cell line, DMEM medium was additionally supplemented with non-essential amino acids (NEAA) (Gibco-11140050). The human monocytic leukemia THP-1 cell line was thawed and cultured in RPMI media supplemented with 20% FBS and 2-mercaptoethanol (Gibco: 21985023) to a final concentration of 0.05 mM at 37°C in a humidified atmosphere at 5% CO_2_. Once the culture was established, the FBS concentration was reduced to 10% for further subculturing. PMA was used at a final concentration of 1000ng/ml for 48 hours to differentiate the THP-1 cells. This was followed by incubating the differentiated THP-1 cell in PMA free media for 24hr. Polarization of the differentiated THP-1 macrophages to M1 phenotype was done by adding 10ng/ml LPS and 50ng/ml IFN-γ (32). For RAW264.7 and THP-1 macrophage intracellular survival assay, 5 x 10^5^ cells were seeded into 12 well plates the day before the experiment. For Caco-2 cells, 5 x 10^5^ cells were seeded into 12 well plates and allowed to grow for one week until tight junctions were formed (49). LMO strains were grown overnight in BHI broth at 37°C. The next day the overnight cultures were inoculated into fresh BHI to a starting OD_600_ of 0.05 and grown for 2 h at 37°C till an OD_600_ of ∼0.2. Cell lines were infected with a MOI of 5. The infection was conducted for 1 hour. The wells were then washed with Dulbecco’s phosphate-buffered saline (PBS) (prewarmed at 37°C), and extracellular bacteria were killed by incubating with DMEM containing 40 µg/ml gentamicin for 1 hour. Medium was changed after 1 hour and intracellular bacteria was harvested by lysing the host cells with 1 ml of ice-cold PBS containing 0.1% Triton X-100. Several serial dilutions were plated on BHI agar plates and kept at 37°C for the bacterial colonies to appear. Colonies were counted manually as well as using the software openCFU (version 3.9.0) (50).

### cDNA preparation and qRT PCR

All RNA extraction was carried out using NucleoSpin RNA Plus kit (Macherey Nagel) according to manufacturer’s protocol. cDNA was synthesized from RNA using the RevertAid reverse transcriptase for first strand cDNA synthesis (ThermoFisher Scientific: K1622). qRT-PCR was performed using SYBR green based iTaq Universal SYBR Green Supermix (Biorad: 1725121), using appropriate primers (Table S2) on a Biorad C1000 thermocycle equipped with CFX96 Optical Reaction Module. The relative expression of various genes was normalized to that of the endogenous reference 16S rRNA gene and the fold change in expression was calculated using the comparative Ct method. For host qRT-PCR, the normalization was done using ACTH as endogenous reference. p-values were calculated using the Two-way ANOVA with Bonferroni’s multiple comparison test.

### Confocal microscopy

To visualize the expression of EutK-mCherry, the recombinant LMO strains LMAC038 (*eutK::mCherry)*, LMAC040 ((*eutK::mCherry*; *ΔeutV*) and LMAC043 (*eutK::mCherry; ΔeutB*), were initially grown overnight in BHI broth at 37°C. Bacterial cells were then washed with sterile PBS and reconstituted in MMWB medium containing either Gln (4.1mM) or EA (16mM) and VitB12 (200µM) as needed and incubated at 37°C on a shaker (180 rpm). Cells were harvested by centrifugation (5000x g) at desired timepoints and were fixed with 4% paraformaldehyde for 10 minutes at 25°C. Cells were then washed thrice with PBS and smear dried on a Poly-L-lysine coated slide (Electron Microscopy Science: 63410), mounted with Prolong Diamond Antifade Mountant (Invitrogen: P36965), covered with #1.5H glass cover slips (Electron Microscopy Sciences), and imaged using the Olympus FLUOVIEW FV3000 confocal microscope equipped with the Fluoview FV315-SW software. Images were acquired in Z-stacks using a step size of 0.20 µm, which were then processed using the Olympus cellSens Dimension software.

Caco-2, RAW264.7 and THP-1 cells were thawed and seeded as mentioned earlier on 6 well plates containing #1.5H glass cover slips (Electron Microscopy Sciences). Cells were grown and maintained as described above. The infection was conducted for 1 hour at an MOI of 5, followed by gentamycin treatment as described above. After infection, the cells were fixed with 4% paraformaldehyde for 10 minutes at 25°C followed by permeabilization with 0.01% Triton X-100 in PBS for 10 minutes at 25°C. The cells were then treated with 5% BSA in PBS for 90 minutes at 25°C followed by treatment with anti-Listeria antibody (Abcam: ab35132) at 25°C for 2 hours. The next day, cells were then washed with PBS and treated with Alexa488 conjugated secondary antibody (Invitrogen: A27034) for 90 minutes at 25°C. Cells were then washed and stained for actin using Alexa Fluor plus 405 Phalloidin (Invitrogen: A30104) for 20 minutes at 25°C. Finally, cells were again washed with PBS, mounted, and imaged as described above.

### Statistical Analysis

Statistical Analyses were done using GraphPad Prism, v9.5.1. P values were calculated using two-way ANOVA with Bonferroni’s multiple comparison test.

### Data Availability

No large datasets were generated that required deposition into public repositories. All data is in the manuscript with the source data underlying the Figures is in Table S3.

## ACKNOWLEDGEMENTS

We thank S. Kolodziej and P. Navarro for sectioning samples for TEM and C. Wu and M.B. Willis for help with strain design and troubleshooting. A special thanks to A. Nagar for advice on culturing and imaging cell lines. Finally, we thank the following members of the LMO community for strains, protocols and advice: D. Portnoy, J.D. Sauer, S. Halbedel and M. O’Riordan. This work was supported by the National Institute of Allergy and Infectious Diseases of the National Institutes of Health under award number R21AI167124 to DAG. The content is solely the responsibility of the authors and does not necessarily represent the official views of the National Institutes of Health. Author contributions: A.C., K.K., and D.A.G. designed research; A.C and K.K performed experiments; A.C and D.A.G analyzed data and wrote the paper.

