## Supplementary material for "Role of Ethanolamine Utilization and Bacterial Microcompartment Formation in *Listeria monocytogenes* Intracellular Infection": Figures S1-S4, Tables S1-S2

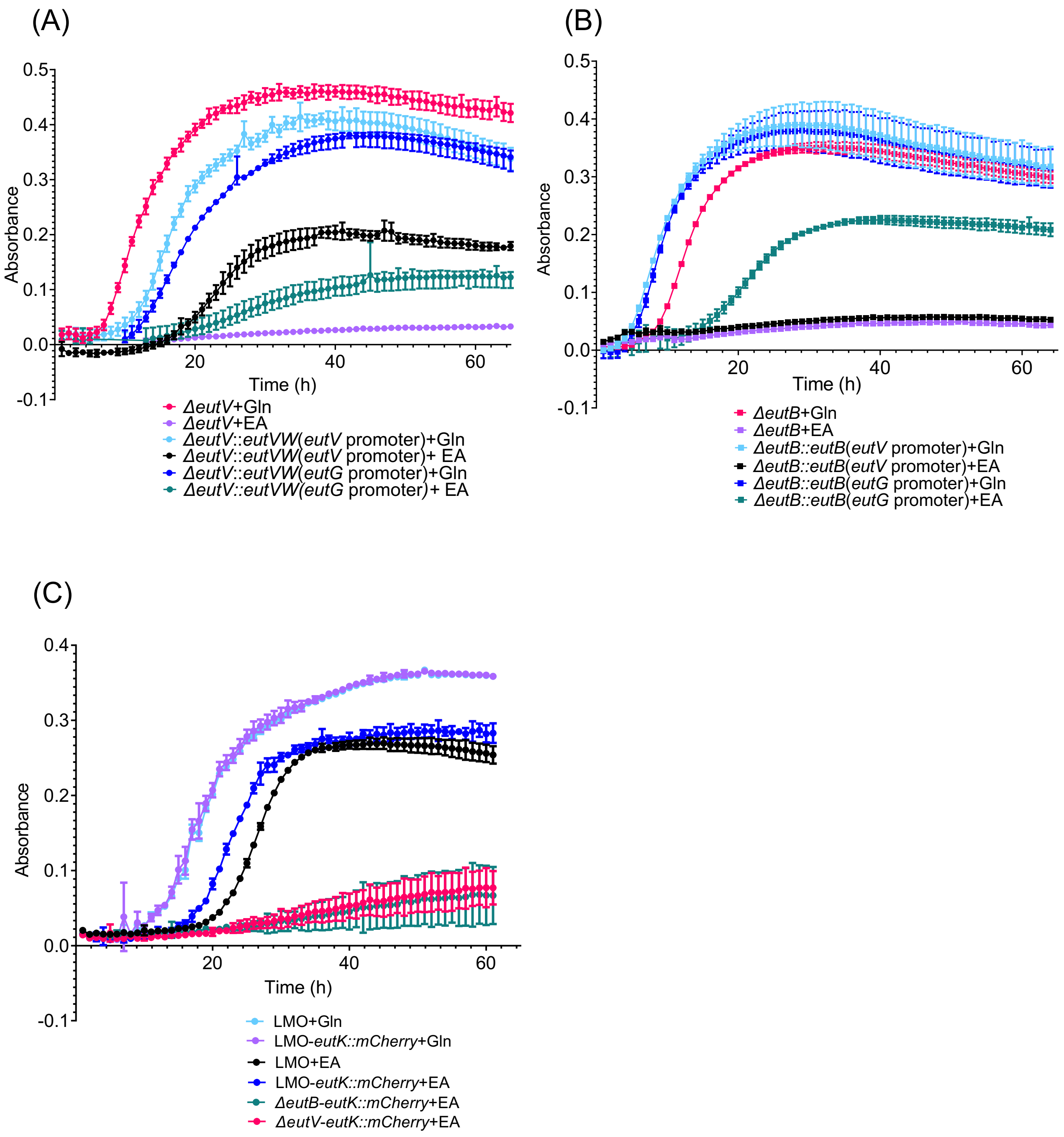

**Fig. S1: Growth curves of engineered strains in MMWB with EA or Gln.** Growth of *ΔeutV* (A) and *ΔeutB* (B) mutants complemented using different *eut* promoters. (C) Growth of *eutK::mCherry.*

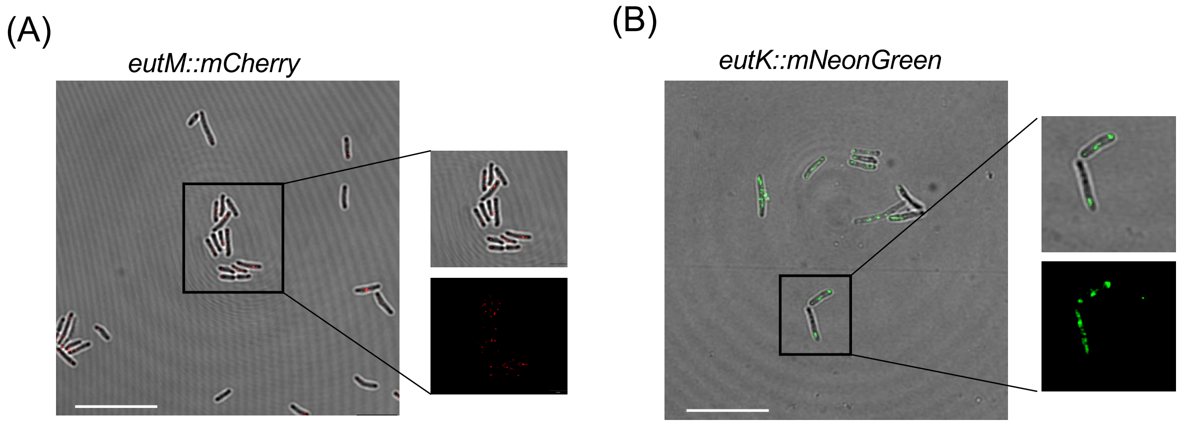

**Fig. S2: EUT BMCs can be fluorescently tagged.** (D) LMO expressing *eutM::mCherry* showed the formation of BMCs in vitro. (E) LMO expressing *eutK::mNeonGreen* also showed the formation of BMCs in vitro. Scale bar represents 10µm.

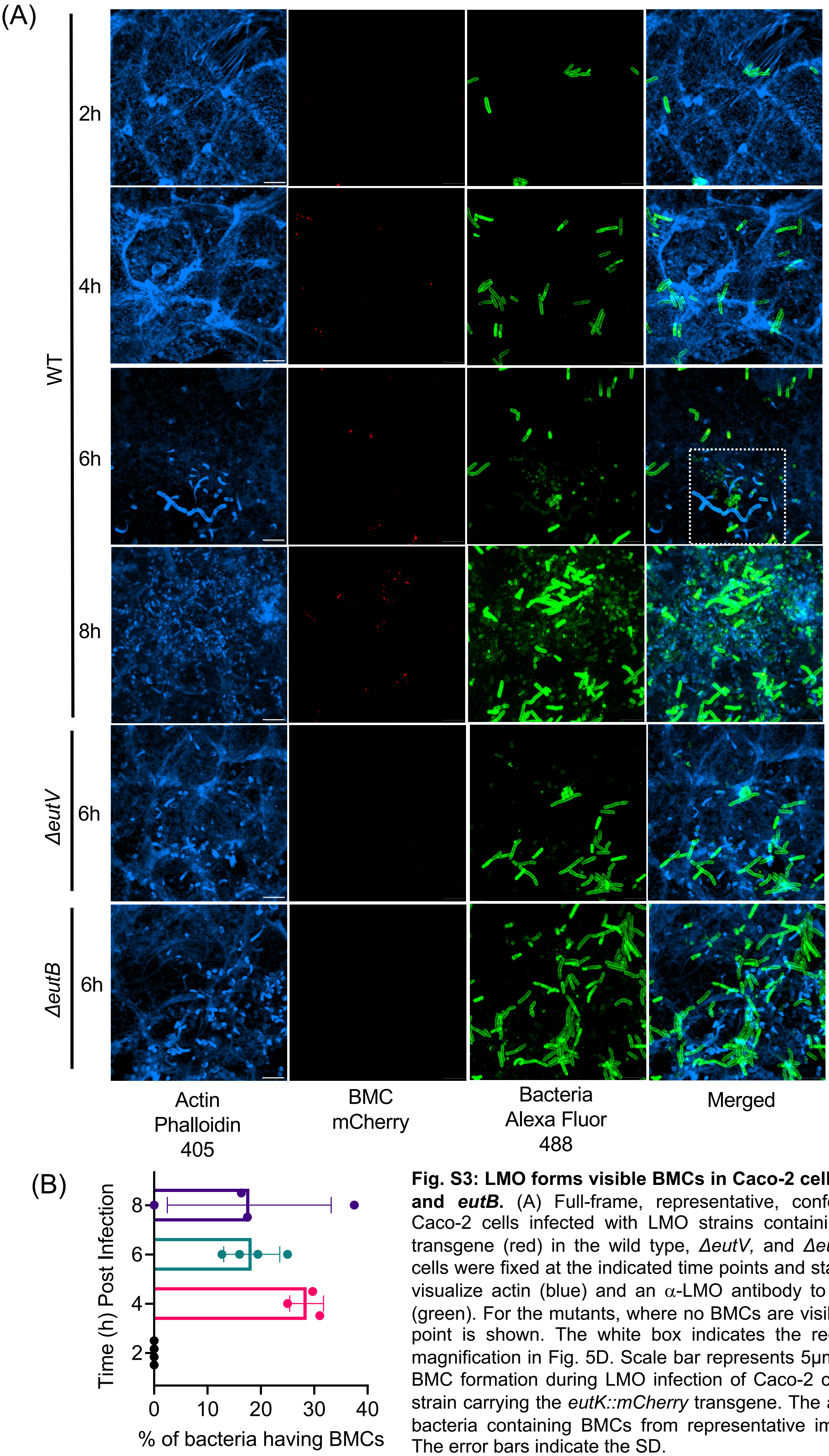

**Fig. S3: LMO forms visible BMCs in Caco-2 cells dependent on *eutV* and *eutB*.** (A) Full-frame, representative, confocal images showing Caco-2 cells infected with LMO strains containing the *eutK::mCherry* transgene (red) in the wild type, *ΔeutV,* and *ΔeutB* backgrounds. The cells were fixed at the indicated time points and stained with phalloidin to visualize actin (blue) and an α-LMO antibody to visualize the bacteria (green). For the mutants, where no BMCs are visible, only a 6-hour time point is shown. The white box indicates the region shown at higher magnification in Fig. 5D. Scale bar represents 5μm. (B) Quantification of BMC formation during LMO infection of Caco-2 cells with the wild type strain carrying the *eutK::mCherry* transgene. The average percentage of bacteria containing BMCs from representative images was calculated. The error bars indicate the SD.

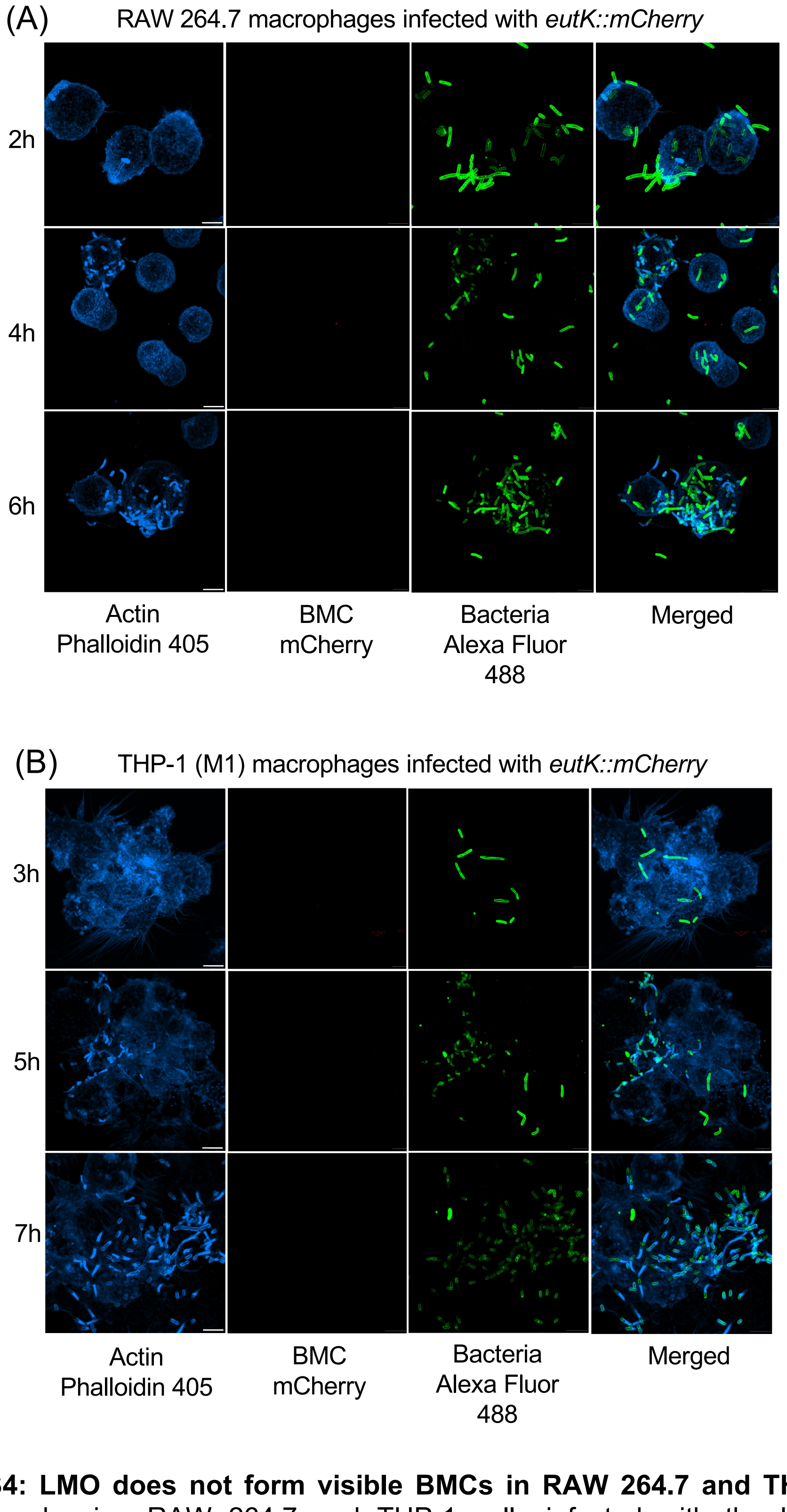

**Fig. S4: LMO does not form visible BMCs in RAW 264.7 and THP-1 cells.** Representative images showing RAW 264.7 and THP-1 cells infected with the LMO strain containing the *eutK::mCherry* transgene (red). The cells were fixed at the indicated time points and stained with phalloidin to visualize actin (blue) and an $\alpha$-LMO antibody to visualize the bacteria (green). Scale bar represents 5μm.

**Table S1: Composition of minimal media (MMWB)**

| Components | Concentration |
|---|---|
| $Na_2HPO4.7H_2O$ | 11.55mM |
| $KH_2PO_4$ | 4.82mM |
| $MgSO_4.7H_2O$ | 1.7mM |
| MOPS (pH 7.4) | 100mM |
| Glucose | 55mM |
| Riboflavin | 1.33μM |
| Thiamine | 2.96 μM |
| Biotin | 2.05 μM |
| Lipoic acid | 24pM |
| Methionine | 0.67mM |
| Cysteine | 0.82mM |
| Leucine | 0.76mM |
| Isoleucine | 0.76mM |
| Valine | 0.85mM |
| Arginine | 0.57mM |
| Histidine | 0.64mM |
| *Glutamine | 4.1mM |
| *Vitamin B12 | 200μM |
| *Ethanolamine | 16mM |

* These were added as required. Both Glutamine and Ethanolamine are used as a nitrogen source. Vitamin B12 was added when Ethanolamine was used as a nitrogen source.

**Table S2: List of bacterial strains and primers used in the study.**

Bacterial strains

| Strain name | Purpose | Comments |
|---|---|---|
| *E. coli* TOP10 | cloning | |
| *E. coli* DH5α | cloning | |
| *Listeria monocytogenes* 10403S (LMO) | Study organism | Source: Bei Resource; wild type |
| LMAC030: *ΔeutV* | Study organism | *ΔeutV* constructed on LMO 10403s background |
| LMAC031: *ΔeutB* | Study organism | *ΔeutB* constructed on LMO 10403s background |
| LMAC007: *eutK::mNeonGreen* | Study organism | LMO expressing *eutK::mNG* |
| LMAC024: *ΔeutB::eutB(eutV promoter)* | Study organism | *ΔeutB::eutB; ΔeutB* complemented with a copy of *eutB* gene under the control of *eutV* promoter. |
| LMAC026: *ΔeutV::eutV(eutG promoter)* | Study organism | *ΔeutV::eutVW; ΔeutV* complemented with a copy of *eutVW* gene under the control of *eutG* promoter. |
| LMAC034: *ΔeutB::eutB(eutG promoter)* | Study organism | *ΔeutB::eutB; ΔeutB* complemented with a copy of *eutB* gene under the control of eutG promoter with riboswitch. |
| LMAC037: *ΔeutV::eutV(eutV promoter)* | Study organism | *ΔeutV::eutVW; ΔeutV* complemented with a copy of *eutVW* gene under the control of *eutV* promoter. |
| LMAC038: *eutK::mCherry* | Study organism | LMO expressing *eutK::mCherry* |
| LMAC039: *eutM::mCherry* | Study organism | LMO expressing *eutM::mCherry* |
| LMAC040: *eutK::mCherry*; *ΔeutV* | Study organism | LMAC030 expressing *eutK::mCherry* |
| LMAC043: *eutK::mCherry*; *ΔeutB* | Study organism | LMAC031 expressing *eutK::mCherry* |

List of primers cloning, knockout (KO) generation, complementation (CT)

| Primer sequence | Purpose | Comment |
|---|---|---|
| TGGGGATCGGAATTCGAGCTGACGGATGCACTGCAACAAATC | *eutV* promoter_fwd | LMAC007: *eutK::mNeonGreen* |
| CATTTGGCATTCTATAGAAGCATTTACTTGAGCGAAAG | *eutV* promoter_rev | |
| CTTCTATAGAATGCCAAATGAAGCGCTTG | *eutK* coding_fwd | |
| AGCTCCCGCTCCTGCGCCGTTTTCACTCCTTCCATAAATATCTTTTAG | *eutK* coding_rev | |
| GGCGCAGGAGCGGGAGCTATGGTGAGCAAGGGCGAGGAGGATA | mNeonGreen_fwd | |
| CGAATTCCTGCAGCCCGGGGTTACTTGTACAGCTCGTCCATGCCCATCAC | mNeonGreen_rev | |
| TGGGGATCGGAATTCGAGCTGACGGATGCACTGCAACAAATC | *eutV* promoter_fwd | LMAC024: *ΔeutB::eutB(eutV promoter)* |
| TTAAAATCATTCTATAGAAGCATTTACTTGAGCGAAAG | *eutV* promoter_rev | |
| CTTCTATAGAATGATTTTAAAAACGAATTTATTCG | *eutB* coding_fwd | |
| CGAATTCCTGCAGCCCGGGGTTATTTTAGGAAAATAGATGCATC | *eutB* coding_rev | |
| TGGGGATCGGAATTCGAGCTAGCAAATTCTTCTTGCAAG | *eutG*Promoter_fwd | LMAC026: *ΔeutV::eutV(eutG promoter)* |
| ACAAAATCATGAGATTACCTCCTAGTTTTTTAAAATTAAAAAAG | *eutG*Promoter_rev | |
| AGGTAATCTCATGATTTTGTCCTATATTGTGAC | *eutVW* coding_fwd | |
| CGAATTCCTGCAGCCCGGGGCTACTTCTTTGTAGCATGG | *eutVW* coding_rev | |
| ACAGATCTATCGATGCATGCAAAAGCAAATGATTACTCGG | *eutV* upstream_fwd | LMAC030: *ΔeutV* |
| ATCATGTTGTTCTATAGAAGCATTTACTTGAG | *eutV* upstream_rev | |
| CTTCTATAGAACAACATGATTCGTAATCTG | *eutV* downstream_fwd | |
| CCTCGCGTCGGGCGATATCGTCATAATAAAGTAATTCCCGG | *eutV* downstream_rev | |

| | | |
|---|---|---|
| ACAGATCTATCGATGCATGCTTAGGTGAAATGGTTGAG | *eutB* upstream_fwd | LMAC031: *ΔeutB* |
| CCATTTTCGCTCGATAAATCCTCCTCTC | *eutB* upstream_rev | |
| GATTTATCGAGCGAAAATGGTAAATTAACTAGC | *eutB* downstream_fwd | |
| CCTCGCGTCGGGCGATATCGAACTGTACGACGAGCTTC | *eutB* downstream_rev | |
| TGGGGATCGGAATTCGAGCTAGCAAATTCTTCTTGCAAG | *eutG* promoter_fwd | LMAC034: *ΔeutB::eutB(eutG promoter)* |
| TTAAAATCATGAGATTACCTCCTAGTTTTTAAAATTAAAAAAG | *eutG* promoter _rev | |
| AGGTAATCTCATGATTTTAAAAACGAATTTATTCG | *eutB* coding_fwd | |
| CGAATTCCTGCAGCCCGGGGTTATTTTAGGAAAATAGATGCATC | *eutB* coding_rev | |
| TGGGGATCGGAATTCGAGCTGACGGATGCACTGCAACAAATC | *eutVW*-promoter+ coding_fwd | LMAC037: *ΔeutV::eutV(eutV promoter)* |
| CGAATTCCTGCAGCCCGGGGCTACTTCTTTGTAGCATGGATAGAATTG | *eutVW*-promoter+ coding_rev | |
| TGGGGATCGGAATTCGAGCTAGCAAATTCTTCTTGCAAG | *eutG* promoter_fwd | AC038 construct. Electroporated to LMO to generate LMAC038: (*eutK::mCherry);* LMAC040 (*eutK::mCherry*; *ΔeutV*) and LMAC043(*eutK::mCherry*; *ΔeutB)* |
| CATTTGGCATGAGATTACCTCCTAGTTTTTAAAATTAAAAAAG | *eutG* promoter_rev | |
| AGGTAATCTCATGCCAAATGAAGCGCTTG | *eutK* coding_fwd | |
| TACTAACCATGTTTTCACTCCTTCCATAAATATCTTTTAG | *eutK* coding_rev | |
| GAGTGAAAACATGGTTAGTAAAGGTGAAG | mCherry coding_fwd | |
| CGAATTCCTGCAGCCCGGGGTTATTTATATAATTCATCCATACCAC | mCherry coding_rev | |
| TGGGGATCGGAATTCGAGCTAGCAAATTCTTCTTGCAAG | *eutG* promoter-fwd | |

| | | |
|---|---|---|
| CGTTTGCCATGAGATTACCTCCTAGTTTTTTAAAATTAAAAAAG | *eutG* promoter-rev | LMAC039: *eutM::mCherry* |
| AGGTAATCTCATGGCAAACGCAAACGCATTAG | *eutM* coding_fwd | |
| TACTAACCATTTCAGCGCTTTTTGGTAGAATTG | *eutM* coding_rev | |
| AAGCGCTGAAATGGTTAGTAAAGGTGAAG | *mCherry* coding_fwd | |
| CGAATTCCTGCAGCCCGGGGTTATTTATATAATTCATCCATACCAC | *mCherry* coding_rev | |

List of primers used for qRT PCR:

| Primer name | Primer sequence |
|---|---|
| ACTBqPCR1S | CACCATTGGCAATGAGCGGTTC |
| ACTTBqPCR1AS | AGGTCTTTGCGGATGTCCACGT |
| IL6qPCR1S | AGACAGCCACTCACCTCTTCAG |
| IL6qPCR1AS | TTCTGCCAGTGCCTCTTTGCTG |
| IL8qPCR1S | GAGAGTGATTGAGAGTGGACCAC |
| IL8qPCR1AS | CACAACCCTCTGCACCCAGTTT |
| IL1bqPCR1S | CCACAGACCTTCCAGGAGAATG |
| IL1bqPCR1AS | GTGCAGTTCAGTGATCGTACAGG |
| ITNFaqPCR1S | CTCTTCTGCCTGCTGCACTTTG |
| ITNFaqPCR1AS | ATGGGCTACAGGCTTGTCACTC |
| CXCL2qPCR1S | GGCAGAAAGCTTGTCTCAACCC |
| CXCL2qPCR1AS | CTCCTTCAGGAACAGCCACCAA |
| MCP1qPCR1S | AGAATCACCAGCAGCAAGTGTCC |
| MCP1qPCR1AS | TCCTGAACCCACTTCTGCTTGG |
| *eutV* qPCR1S | TGTTGTAGGGGAAGCGACAG |
| *eutV* qPCR1AS | GATAATTCCGCCAGCAAGCC |
| *eutA* qPCR1S | GGAGGCGTTGCAGACTGTATT |
| *eutA* qPCR1AS | CGACCACTGTTGCCCGTAT |
| *eutB* qPCR1S | GCTCATTTCGGCGTAGACCA |
| *eutB* qPCR1AS | CCGGCGCGAATTACTTGTTT |
| *eutK* qPCR1S | GCTTTCTTGGTGCCGTTGTT |
| *eutK* qPCR1AS | TTCAGTTGCCGCTTTTCCTG |
| *eutM* qPCR1S | CAAGTTGGTGGCGGTCTAGT |
| *eutM* qPCR1AS | TTGCGTCTACTTCGCTGTGT |
| 16s qPCR1S | GGGGCTAATACCGAATGATAGAATG |
| 16s qPCR1AS | TATGCATCGTTGCCTTGGTAG |
| *pduA* qPCR1S | TCGGGTCTGGACTGGTTACT |
| *pduA* qPCR1AS | CCACATCAGTATGCGGACGA |
| *pduQ* qPCR1S | CAGGCAAAATCCAACCGCAA |
| *pduQ* qPCR1AS | GCTGTTACTTCCGAGCCTGT |
